# Prot2Surf: fast analysis of protein–surface binding modes

**DOI:** 10.64898/2026.08.26.747352

**Authors:** Abraham Muñiz-Chicharro, Gamze Tanriver, Artur Góra

**Author notes:** These authors contributed equally to this work (Abraham Muñiz-Chicharro, Gamze Tanriver).

## Abstract

**Summary:** Prot2Surf is a software tool designed for the characterization and prediction of protein association to surfaces. In this application note, Prot2Surf was tested using catalytic domains of the lytic polysaccharide monooxygenases (LPMOs), interacting with native surfaces. The results show that the software can efficiently analyze key binding features, including protein–surface distances, distances between catalytically reactive atoms, and the orientation angle between surface chains and the protein. These features are essential for distinguishing productive binding poses in these protein–surface systems and for understanding interaction patterns that provide guidance on protein engineering. Prot2Surf performs these analyses within seconds to a few minutes, providing a fast and accessible framework to post-process and characterize protein-surface encounter complexes.

**Availability and implementation:** Prot2Surf, which is written in Fortran90, is documented and freely available as open source on GitHub: https://github.com/TUNNELING-GROUP/Prot2Surf. In order to run Prot2Surf, users should also install the SDA software package which is freely available at https://www.h-its.org/downloads/sda7/.

## 1 Introduction

Experimental characterization of protein–surface association features, such as binding modes and affinities, remains challenging. A limited amount of crystal structures of protein-surface complexes is available. (Frandsen *et al*. 2016, 2017, Bissaro *et al*. 2018, Ciano *et al*. 2020, Tandrup *et al*. 2020, 2022). Moreover, only partial segments of surface, such as a few single oligosaccharide chains, are resolved, often forming limited or no contacts with the binding surface of protein. Since there are no nearby chains and surface layers that affect association and orientation of protein on surface, the adequate description of the possible binding modes and interactions of protein on surface is not accessible. Therefore, modelling and prediction of protein association on surfaces are crucial for elucidating the molecular mechanisms underlying protein–surface binding, ultimately enabling rational protein design to increase affinity for the target surface. Current computational strategies are largely constrained by computational costs. For instance, fully atomistic explicit-solvent molecular dynamics (MD) simulations are highly demanding, and with current computational resources, only a few microseconds are typically simulated within days for protein–surface systems, due to their large size. Since protein–surface association is largely diffusion-driven, capturing the full diffusional process and identifying encounter complexes may require simulations reaching millisecond timescales, making exhaustive MD simulations computationally unfeasible. Brownian dynamics (BD) simulations provide an efficient alternative, as they reduce computational cost by using approximations, such as rigid-body representations and implicit-solvent models while retaining the essential features of long-range diffusional association. Among the available software, SDA7 (Gabdoulline R. R. and Wade 1997, Gabdoulline Razif R. and Wade 1998, Martinez *et al*. 2015) stands out by offering several types of simulations: protein-protein, protein-ligand, protein-surface and multiple molecule simulations. However, this software does not provide a clustering tool to fully characterize protein–surface complexes. To address this limitation, we developed Prot2Surf, a software tool, to analyze different features of encounter complexes: distance to surface, distance between reactive atoms and angular alignment between the active site (AS) of the protein and the surface chain. In the content of this work, we also evaluated the performance of Prot2Surf with LPMO–cellulose and LPMO– chitin systems, demonstrating its efficiency and accuracy on detecting productive binding modes and mapping surface binding residues that contribute to the association.

## 2 Implementation

### 2.1 Overview

The Prot2Surf software tool was developed to post-process rigid-body encounter complexes produced by the SDA7 software (Gabdoulline R. R. and Wade 1997, Gabdoulline Razif R. and Wade 1998, Martinez *et al*. 2015). Prot2Surf reads the SDA7 encounter complexes files, which contain one translation vector and two rotation vectors for each encounter complex. These vectors are used to reconstruct the solute coordinates by applying the corresponding rigid-body transformation to the mobile solute PDB used in the SDA7 simulations. The clustering selection protocol utilizes Near Attack Configuration (NAC) concept, in a way that encounters complexes are compared based on defined geometrical parameters: distance and angle, providing access to rational classification of encounters. The reconstruction uses the reference centers of the surface and protein reported in the encounter-complexes header. As for the software architecture, this is designed around a modular workflow to (i) compute distance arrays between the encounters and the surface, (ii) cluster encounters based on these distance arrays using an average-linkage hierarchical scheme, and (iii) perform additional analyses such as threshold-based filtering and residue–surface proximity screening. The advantage of the software is that it is not necessary to form a distance matrix since the descriptors to be compared are between the encounter complexes and the surfaces, and not within the encounter complexes. This strategy drastically reduces computational costs. Moreover, OpenMP parallelization is used to the computationally intensive loops over encounters, and pairwise comparisons are parallelized using OpenMP to exploit shared-memory multicore CPUs.

### 2.2 Installation, input, and workflow

The software is written in Fortran and is compiled using make. A Makefile is provided with the source code, so the program can be compiled directly from the command line using the make all command. For compilation, it is necessary to have gfortran installed, together with the required BLAS and LAPACK libraries.

Four main modules are generated during compilation: make_data, clust, threshold, and contact_map. The make_data module reads the SDA7 encounter complexes files to generate the data array required by the clust and threshold modules. The clust module uses this array for clustering, while the threshold module uses it for threshold filtering. Moreover, the output generated by clust and threshold may serve as input for the make_data and contact_map modules. However, the output generated by contact_map cannot be provided as input to any other module (Figure 1). The software also provides auxiliary python script tools to aid in the automatization of tasks such as plotting or regioselectivity calculation.

**Figure 1.**
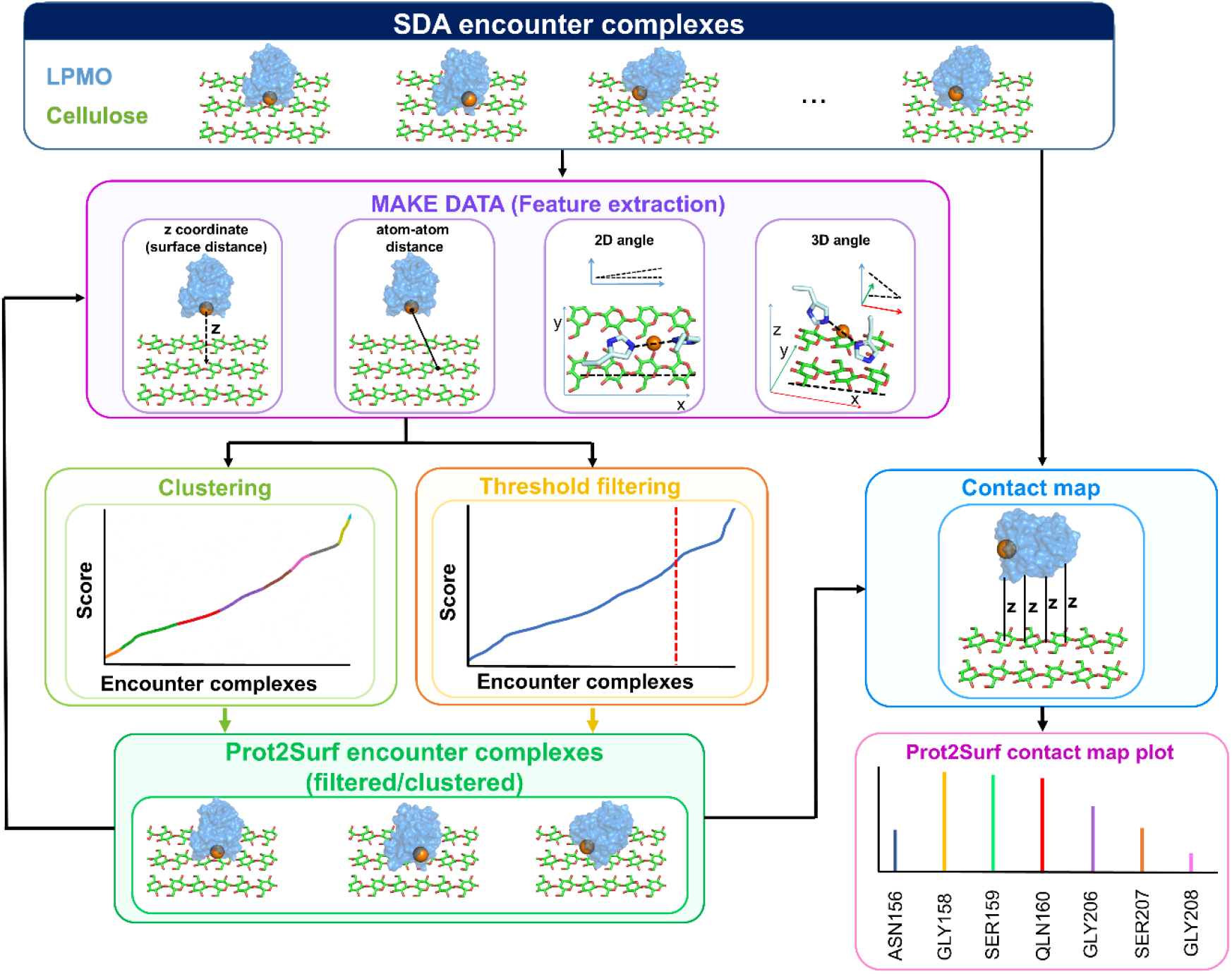
Schematic overview of Prot2Surf workflow.

### 2.3 Output and visualization

After running the clust module, separate encounter-complex files are generated, each containing the encounters assigned to the different clusters, thus files containing information of the different clusters: cluster population, cluster mean, and cluster standard deviation. When the threshold module is used, a single encounter-complex file is generated, containing only the encounters that satisfy the imposed threshold. If contact_map is run, a DAT file is generated containing the number of residues among all encounter complexes whose center-of-mass z coordinate is below a user-defined threshold, set to 6 Å by default.

## 3 Application

Prot2Surf systematically scrutinizes the diffusional encounter complexes of protein associations based on distances and angles. As a particular example, monocopper metalloenzymes Lytic Polysaccharide Monooxygenases which have a conserved histidine-brace motif that coordinates a copper ion and a flat binding surface were selected. In LPMO-native surface systems, initiation of the reaction and the consequent oxidation at the C1 carbon, C4 carbon, or mixed C1/C4 positions, leading to glycosidic bond cleavage, depend on the productive positioning and orientation of surface chains with respect to the Cu–His brace active site of LPMOs.(Walton and Davies 2016, Vaaje-Kolstad *et al*. 2017, Rovira *et al*. 2020, Zhou and Zhu 2020) Therefore, as NAC parameters the copper surface distance and Cu-His-C1-C4 angle need to be evaluated, which are crucial to properly investigate the productive complexes association. In the case study presented in section 3.1, Brownian dynamics simulations were performed using SDA7 (version 7.3.5) (SDA protocol available in the Supplementary Information). Afterwards, we developed a protocol with Prot2Surf that first uses the make_data and clust module to cluster encounter complexes based on the z-coordinate of the copper ion (Step 1). In this analysis, copper z-coordinates close to 0 were interpreted as indicating proximity to, and potential interaction with, the surface. The first cluster, corresponding to the copper ions closest to the surface, was filtered using a 7.5 Å distance threshold to either the C1 or C4 reactive atom (Step 2), using the make_data and threshold modules. After the filtering, 2D-angle clustering between the histidine brace nitrogen of LPMO coordinating the copper ion and surface vectors were performed with make_data and 2D_angle modules to assess productive positions (i.e.: parallel orientations) (Step 3). Then, the first and last clusters of angle analysis were analyzed and classified as C1, C4 or mixed C1/C4 in terms of regioselectivity using the auxiliary tools (Step 4). Lastly, the residues in direct contact with the surface were identified with a threshold distance of residue center of geometry z coordinate below 6.0 Å to map LPMO–surface interactions (Step 5). Note that as illustrated in Figure 1 Prot2Surf provides flexibility to adjust various feature extractions and clustering parameters/order for enzymes with different ASs and differently defined NAC.

### 3.1 Example cases

Prot2Surf was tested on LPMO-surface systems to assess the efficacy and performance of the software and to demonstrate its applicability to protein–native surface systems. The reason behind LPMO selection is their crucial role in depolymerization of recalcitrant polysaccharides *via* the oxidative cleavage, making them to exhibit unique catalytic activities with growing interest in the scientific and industrial fields(Tandrup *et al*. 2018, Forsberg *et al*. 2019). Two LPMOs were selected from different clades, namely the chitin-active SmAA10A and the cellulose-active NcAA9C. It should be noted that while SmAA10A only includes a catalytic domain (CD), native NcAA9C enzyme contains both a CD and a Carbohydrate-Binding Module (CBM) which is connected to CD *via* a flexible linker. In this study, only CDs of the LPMOs were tested. Additionally, the surfaces used are cellulose Iβ and β-chitin, two abundant polysaccharides that are common in nature. In Figure 2A, Cluster 1, which contains the encounter complexes with the smallest copper-ion z-coordinates, indicates that the active sites of SmAA10A and NcAA9C bind to the chitin and cellulose surfaces, respectively. As for productive orientation, Figure 2B shows that SmAA10A is populated in two clusters and many ASs are oriented on the chain in the range of 0-25°. In contrast, the NcAA9C case shows a greater variety of clustering tendencies with a broad range of protein orientation on the surface. The higher number of encounter complexes within 0-25° suggests a higher affinity of SmAA10A towards chitin compared to NcAA9C towards cellulose. The difference in binding affinity observed for NcAA9C may be attributed to the removal of its CBM, which facilitates/mediates binding of enzymes. This result aligns with the observation of previous experimental studies (Kracher *et al*. 2018, Støpamo *et al*. 2024) and suggests the sensitivity and applicability of the combined SDA and Prot2Surf analysis. By analyzing the angular alignment between the LPMO histidine brace and the surface, Prot2Surf provides data that are difficult to obtain experimentally.

**Figure 2.**
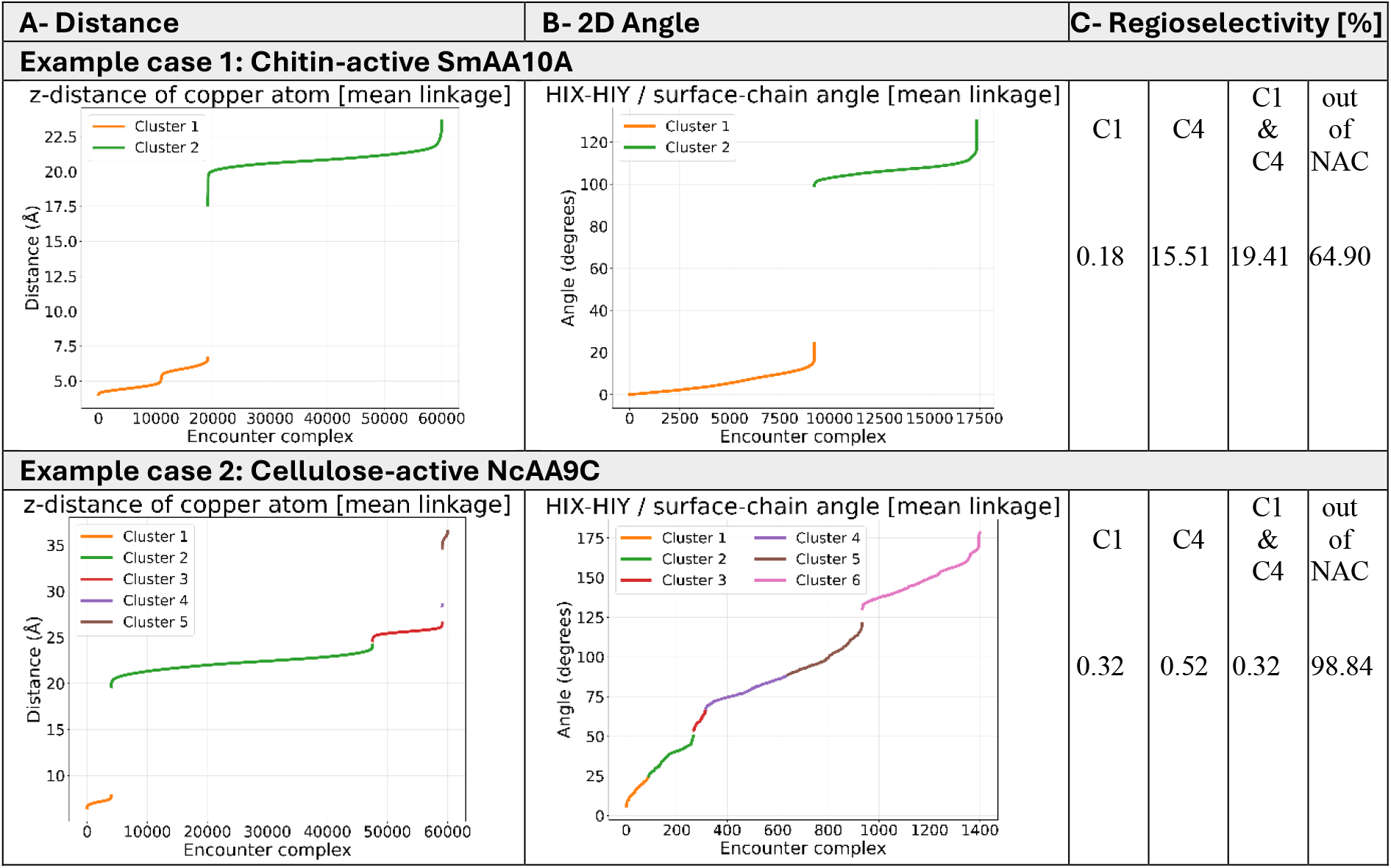
A) Z-coordinate cluster analysis of copper ion, B) 2D angle cluster analysis, C) Analysis of the fulfillment of the NAC condition (%). Raw output data are available in Table S1.

After clustering and filtering the encounter complexes, the regioselectivity was calculated based on the number of occurrences for encounter complex whose histidine brace was aligned with the surface chain and whose copper ion was within 7.5 Å to either C1, C4 or both reactive atoms (Figure 2C). SmAA10A and NcAA9C are C1-oxidizing and C4-oxidizing, respectively. The amount of encounter complexes showing C4 regioselectivity is higher than C1 or mixed C1/C4 for NcAA9A, which is in line with experimental studies.(Borisova *et al*. 2015, Støpamo *et al*. 2024) Although the occurrence of C1 regioselective positions of SmAA10A is low, it is possible that the mixed C1/C4 regioselectivity arises from inaccuracies of the rigid body model. To overcome this issue, the results of Prot2Surf can be used as a starting point for further refinement through MD simulations, enabling analysis of other important factors influencing regioselectivity. Lastly, the differences in the residue contact plots for the two LPMOs in Figure 3 show that the encounter complexes adopt clearly distinct orientations across the different clusters. Residues forming the binding site are frequently involved in close surface contacts in cluster 1, where the copper atom is closest to the surface (Figure S4). Conversely, clusters formed by larger copper–surface distances display more diverse and less defined orientations, a pattern that is particularly evident in cluster 2 of SmAA10A. The outcomes of this analysis provide insight into plausible surface-binding residues by paving the way for future protein design.

**Figure 3.**
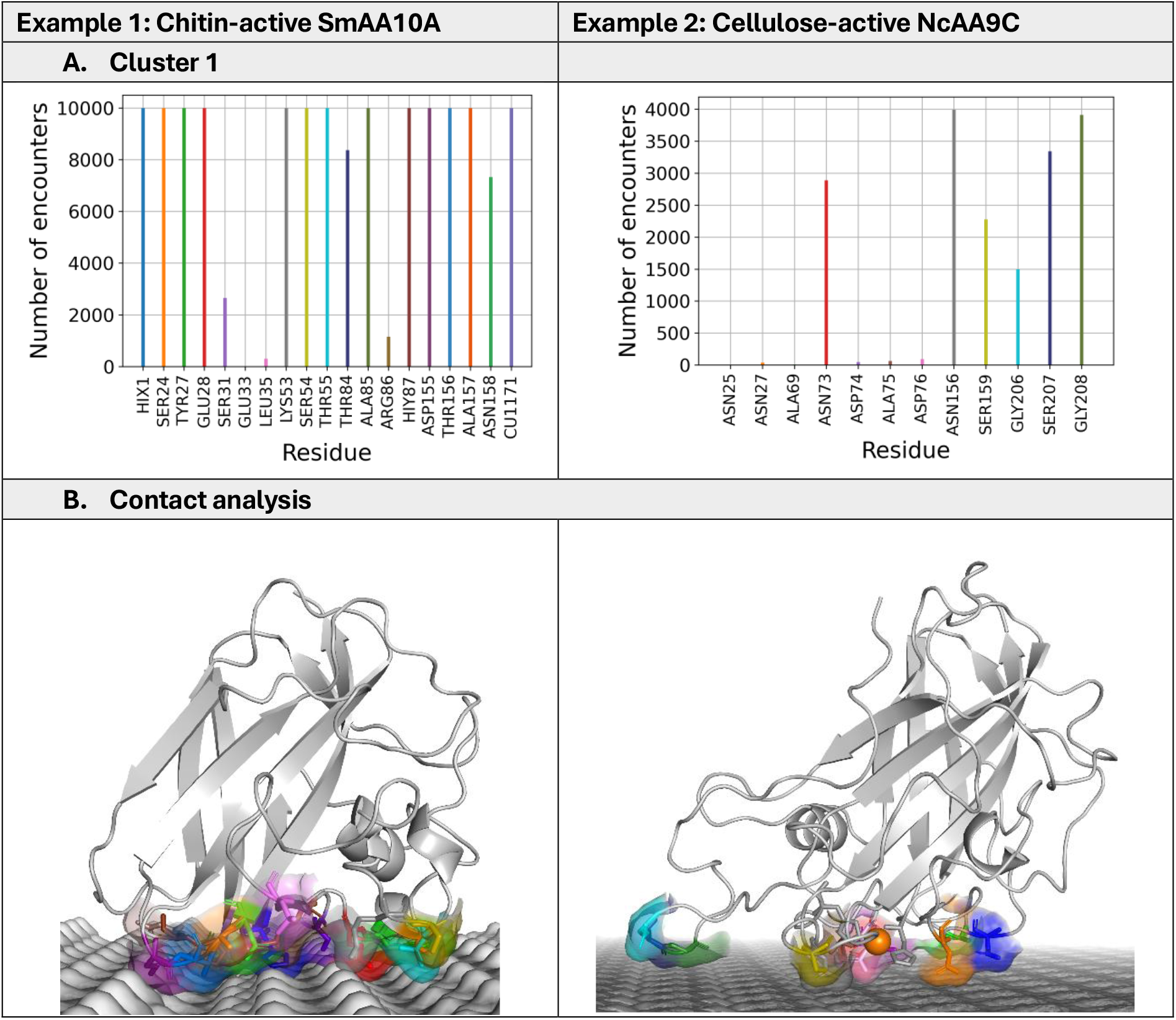
**A**. Close contact residues *vs* number of encounter complexes for the first clusters. **B**. The surface and stick presentations of surface contact residues. Contact residues were colored according to the colors of their corresponding vertical lines in Figure 3A.

## 4 Conclusions

We developed Prot2Surf, a Fortran-based software package that uses mean-linkage clustering and threshold filtering for the prediction and characterization of protein association on surfaces. The program has been tested on LPMOs on cellulose and chitin by combining distance- and angle–based clustering with a threshold filtering and a residue-level contact analysis. The evaluation of the example cases indicates that Prot2Surf allows for the efficient stepwise identification of binding orientations, classification of interaction patterns, and mapping of the closest contacts, thereby facilitating the prediction protein-surface association. Moreover, due to its fast characterization of encounter complexes, Prot2Surf enables the possibility for future studies towards protein design with a focus on improving affinity and/or changing the regioselectivity to desired surfaces by looping over mutagenesis in silico. Furthermore, Prot2Surf is an open-source and user-friendly program with future perspectives on post-processing the outputs of other BD software such as Browndye or GeomBD3.

## Acknowledgements

We would like to thank Rebecca C. Wade for her valuable advice and Karolina Mutisinska, Katarzyna Szleper-Shendy, and Michał Chyski for the preliminary trials.

## Author contributions

Abraham Muñiz-Chicharro (Conceptualization [equal], Data curation [equal], Formal analysis [equal], Investigation [equal], Methodology [lead], Project administration [equal], Software [lead], Validation [equal], Visualization [equal], Writing—original draft [equal], Writing—review & editing [equal])

Gamze Tanriver (Conceptualization [equal], Data curation [equal], Formal analysis [equal], Investigation [equal], Methodology [lead], Project administration [equal], Software [supporting], Validation [equal], Visualization [equal], Writing—original draft [equal], Writing— review & editing [equal])

Artur Gora (Conceptualization [lead], Funding acquisition [lead], Data curation [supporting], Project administration [supporting], Resources [lead], Supervision [lead], Writing—review & editing [supporting])

## Data availability

Code and documentation are available at https://github.com/TUNNELING-GROUP/Prot2Surf. The example cases dataset of this study is available on Zenodo: https://zenodo.org/records/22003489.

## References

Bissaro B, Isaksen I, Vaaje-Kolstad G et al. How a Lytic Polysaccharide Monooxygenase Binds Crystalline Chitin. Biochemistry 2018;57(12):1893–906. 10.1021/acs.biochem.8b00138.

Borisova AS, Isaksen T, Dimarogona M et al. Structural and functional characterization of a lytic polysaccharide monooxygenase with broad substrate specificity. Journal of Biological Chemistry 2015;290(38):22955–69. 10.1074/jbc.M115.660183.

Ciano L, Paradisi A, Hemsworth GR et al. Insights from semi-oriented EPR spectroscopy studies into the interaction of lytic polysaccharide monooxygenases with cellulose. Dalton Transactions 2020;49(11):3413–22. 10.1039/C9DT04065J.

Forsberg Z, Sørlie M, Petrović D et al. Polysaccharide degradation by lytic polysaccharide monooxygenases. Curr Opin Struct Biol 2019;59:54–64. 10.1016/j.sbi.2019.02.015.

Frandsen KEH, Poulsen JCN, Tandrup T et al. Unliganded and substrate bound structures of the cellooligosaccharide active lytic polysaccharide monooxygenase LsAA9A at low pH. Carbohydr Res 2017;448(6001):187–90. 10.1016/j.carres.2017.03.010.

Frandsen KEH, Simmons TJ, Dupree P et al. The molecular basis of polysaccharide cleavage by lytic polysaccharide monooxygenases. Nat Chem Biol 2016;12(4):298–303. 10.1038/nchembio.2029.

Gabdoulline R. R., Wade RC. Simulation of the diffusional association of barnase and barstar. Biophys J 1997;72(5):1917–29. 10.1016/S0006-3495(97)78838-6.

Gabdoulline Razif R., Wade RC. Brownian Dynamics Simulation of Protein–Protein Diffusional Encounter. Methods 1998;14(3):329–41. 10.1006/METH.1998.0588.

Kracher D, Andlar M, Furtmüller PG et al. Active-site copper reduction promotes substrate binding of fungal lytic polysaccharide monooxygenase and reduces stability. Journal of Biological Chemistry 2018;293(5):1676–87. 10.1074/jbc.RA117.000109.

Martinez M, Bruce NJ, Romanowska J et al. SDA7: A modular and parallel implementation of the simulation of diffusional association software. J Comput Chem 2015;36(21):1631–45. 10.1002/jcc.23971.

Rovira C, Walton PH, Wang B et al. Activation of O2 and H2O2 by lytic polysaccharide monooxygenases. ACS Catal 2020;10(21):12760–9. 10.1021/acscatal.0c02914.

Støpamo FG, Sulaeva I, Budischowsky D et al. The Impact of the Carbohydrate-Binding Module on How a Lytic Polysaccharide Monooxygenase Modifies Cellulose Fibers. published online 2024. 10.1186/s13068-024-02564-8.

Tandrup T, Frandsen KEH, Johansen KS et al. Recent insights into lytic polysaccharide monooxygenases (LPMOs). Biochem Soc Trans 2018;46(6):1431–47. 10.1042/BST20170549.

Tandrup T, Muderspach SJ, Banerjee S et al. Changes in active-site geometry on X-ray photoreduction of a lytic polysaccharide monooxygenase active-site copper and saccharide binding. IUCrJ 2022;9(Pt 5):666–81. 10.1107/S2052252522007175.

Tandrup T, Tryfona T, Frandsen KEH et al. Oligosaccharide Binding and Thermostability of Two Related AA9 Lytic Polysaccharide Monooxygenases. Biochemistry 2020;59(36):3347–58. 10.1021/ACS.BIOCHEM.0C00312.

Zhou X, Zhu H. Current understanding of substrate specificity and regioselectivity of LPMOs. Bioresour Bioprocess 2020;7(1):11. 10.1186/s40643-020-0300-6.

