## Supplementary information for "Prot2Surf: fast analysis of protein–surface binding modes"

This Supplementary Information expands the theory and application behind the software and provides all technical details to run Prot2Surf efficiently.

### Distance threshold definition

The threshold definition used to stop the clustering process is defined as:

$$\text{threshold} = \alpha * \text{mean} - (1 - \alpha) * \text{std}$$

Where std is the standard deviation, and $\alpha$ is a normalized value of the standard deviation defined as:

$$\alpha = \frac{\text{std}}{\sqrt{\text{mean}^2 \times \text{std}^2}}$$

This definition of the threshold partially adapts to the distribution of the data. When the standard deviation is low, the threshold takes a small value, which causes clustering to stop quickly because the data are already tightly compacted. When the standard deviation is high, the threshold approaches the mean value, which prevents merging clusters that represent data with substantial differences (Figure S1).

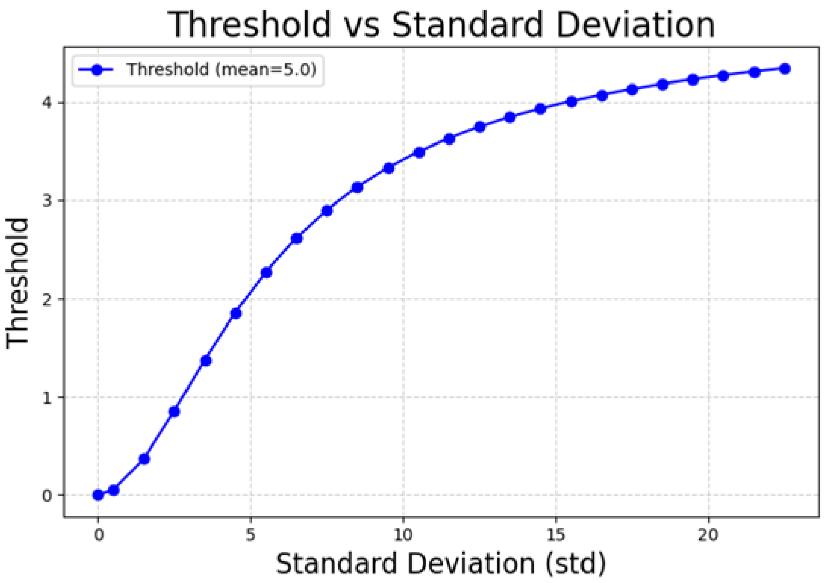

**Figure S1.** The threshold function is based on the mean distance and standard deviation of the matrix distance used in clustering steps.

However, although the mean and standard deviation provide useful global information, they may not accurately represent the central region of asymmetric distributions. Therefore, the middle_range value is compared with the reference value mean_dist+2·standard_deviation to evaluate whether the spread of the distribution is consistent with its central range. This comparison is introduced through the correction factor:

$$\text{factor} = 1 - \frac{|\text{middle\_range} - (\text{mean}_{\text{dist}} + 2 \cdot \text{standard\_deviation})|}{\text{middle\_range} + \text{mean}_{\text{dist}} + 2 \cdot \text{standard\_deviation}}$$

This factor is then used to correct the original clustering distance threshold:

$$\text{threshold}_{\text{corrected}} = \frac{\text{threshold}}{\text{factor}}$$

When middle_range and mean_dist+2· standard_deviation are similar, the numerator of the correction factor is small, and factor remains close to 1. As a result, dividing threshold by this factor barely modifies the clustering threshold. This occurs in Distribution 1 (Figure S2), where the standard deviation is low, and also in Distribution 2 (Figure S3), where the standard deviation is higher, but the reference value remains close to the middle range.

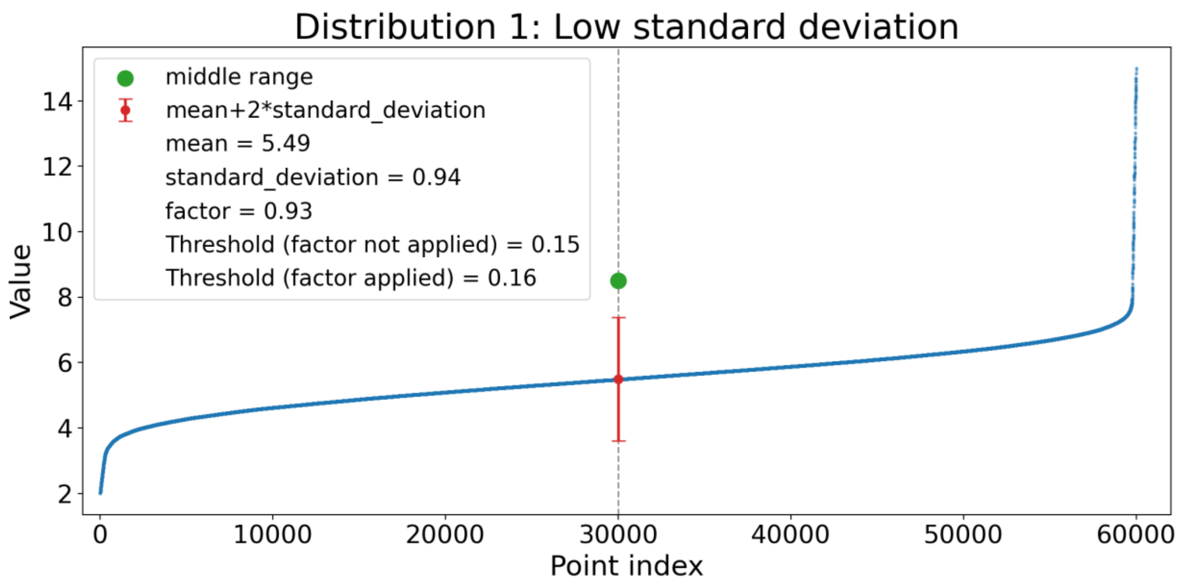

**Figure S2.** Distribution data representing low standard deviation.

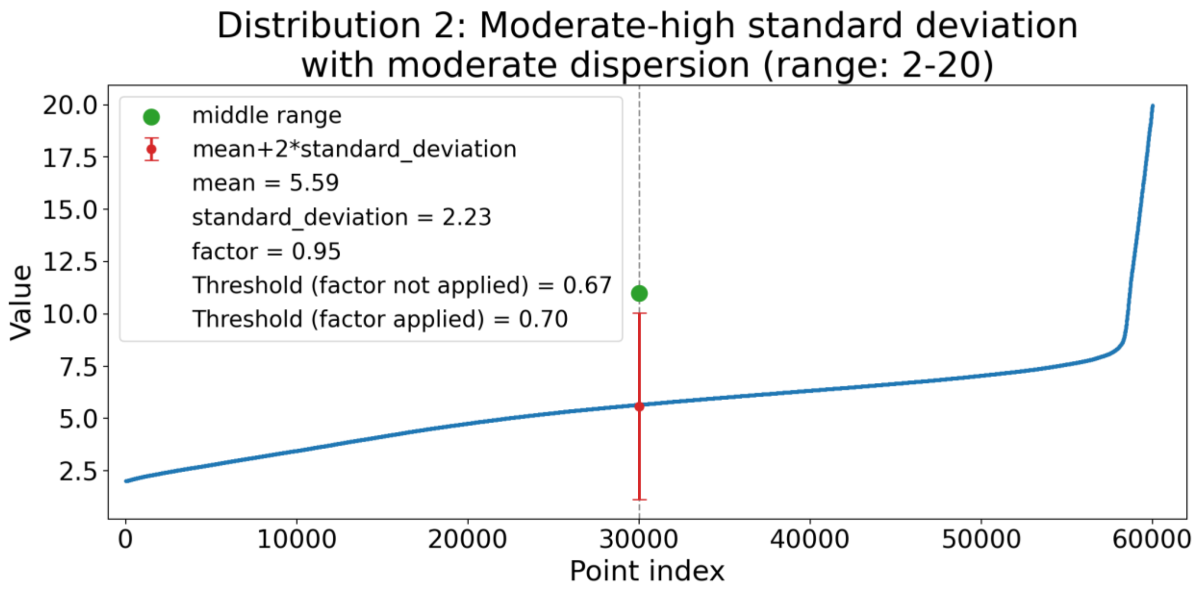

**Figure S3.** Distribution data representing high standard deviation with moderate data dispersion.

In contrast, Distribution 3 represents a case in which the standard deviation is high and mean_dist+2 · standard_deviation differs substantially from middle_range. This increases the numerator of the correction factor, causing factor to move away from 1. Since the original dist_threshold is divided by this factor, a smaller factor produces a larger corrected threshold.

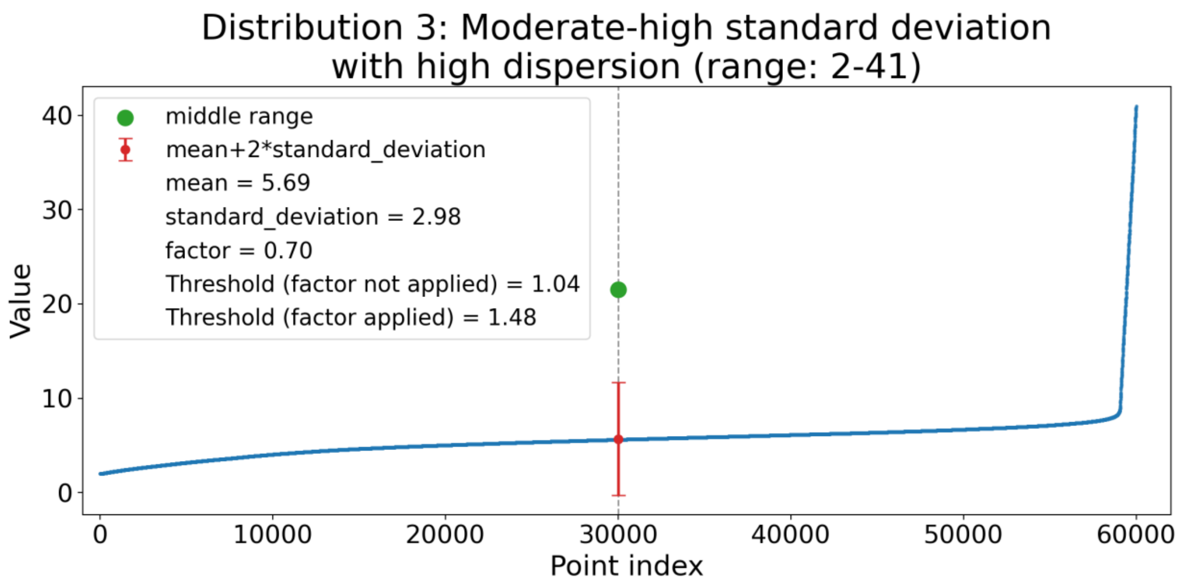

**Figure S4.** Distribution data representing high standard deviation with high data dispersion.

This correction is important because it makes the clustering criterion more conservative when the data are highly dispersed. In such cases, stopping the clustering process too early could artificially split points that still belong to the same broader cluster. By increasing the original distance threshold only when the statistical reference value is not aligned with the middle range, the method accounts for data variability while preventing the clustering decision from being controlled only by the mean and standard deviation.

### User guide

This section provides a practical guide for installing and using Prot2Surf, including installation requirements, supported computing environments, and representative examples.

### Installation and dependencies

To install and run Prot2Surf:

```
git clone https://github.com/TUNNELING-GROUP/Prot2Surf
```

Build dependencies for the Fortran executables:

- gfortran >= 9.0, with OpenMP support through the -fopenmp flag
- OpenMP >= 4.5, provided by the gfortran runtime
- LAPACK and BLAS libraries, linked with -llapack and -lblas
- GNU Make

Optional dependencies for documentation generation:

- Doxygen
- Graphviz

- texlive-latex-base
- texlive-latex-extra
- texlive-fonts-recommended

Dependencies for the auxiliary Python tools in the tools/ directory:

- Python >= 3.8
- numpy
- Matplotlib

On Debian/Ubuntu systems, the required build dependencies can be installed with:

```
sudo apt-get install -y gfortran liblapack-dev libblas-dev make \
   python3 python3-pip
```

The required Python packages can then be installed with:

```
pip install numpy matplotlib
```

#### Computing environments and platforms

Prot2Surf has been tested on Linux-based systems, including Ubuntu 22.04 on x86_64 architecture and high-performance computing environments using the SLURM job scheduler.

The minimum hardware requirements are modest. Any modern x86_64 processor can be used, although multi-core CPUs are recommended when using OpenMP parallelization. Memory requirements depend on the number of encounter complexes and on the type of analysis performed.

For array-based clustering of 60,000 encounter complexes, the approximate memory usage is below 65 MB. Therefore, a standard workstation with 4 GB of RAM is sufficient for this type of analysis.

#### Examples

The main Prot2Surf workflow is composed of four executables:

1. make_data computes encounter-derived numerical arrays.
2. threshold filters encounter complexes according to a numerical cutoff.
3. clust performs hierarchical clustering on the selected data.
4. contact_map analyzes residue proximity across a set of encounter complexes.

Each executable can be invoked with the -help option to display the complete list of available arguments. All examples below assume that the compiled binaries are available in the bin/ directory and that the input files are located in the working directory.

#### Example 1. Computing encounter-derived arrays with make_data

The make_data executable generates a numerical array from the encounter complexes. The type of data to be computed is selected with the -data_type option. Supported data types include rmsd, z_coord, atoms_dist, 2D_angle, and 3D_angle.

```
./make_data \
   -pdb2       p2_noh.pdb      \
   -atoms2     Cu              \
   -complexes  assoc_complexes \
   -data_type  z_coord         \
   -output     array_z.txt
```

```
./make_data \
   -pdb1       p1_noh.pdb      \
   -atoms1     C1              \
   -pdb2       p2_noh.pdb      \
   -atoms2     Cu              \
   -complexes  assoc_complexes \
   -data_type  atoms_dist      \
   -output     array_dist.txt
```

The expected output is a text file, such as array_z.txt or array_dist.txt, containing one numerical value per encounter complex. This file can be used directly as input for threshold or clust.

#### Example 2. Filtering encounter complexes with threshold

The threshold executable reads a numerical array generated by make_data, applies a user-defined cutoff, and writes both the filtered array and the corresponding filtered encounter-complex file.

```
./threshold \
   -array_input     array_C1.txt              \
   -complexes       Cu_z_cluster0001_complexes \
   -array           threshold_C1_array.txt    \
   -complexes_output threshold_C1_complexes   \
   -cutoff          5.0
```

This command generates threshold_C1_array.txt, containing the array values for the selected encounters, and threshold_C1_complexes, containing the filtered encounter complexes.

#### Example 3. Hierarchical clustering with clust

The clust executable performs hierarchical clustering using minimum, maximum, or mean linkage. The input can be either a one-dimensional array or a two-dimensional distance matrix.

```
./clust \
   -complexes     assoc_complexes        \
```

```
  -input          Cu_z_array.txt          \
  -nb_encounters  5000                     \
  -output_name    Cu_z                     \
  -linkage        mean
```

The expected output files are Cu_z_clust_info.txt, Cu_z_clusters.txt, and one association file per cluster, named Cu_z_cluster*_complexes.

#### Example 4. Residue proximity analysis with contact_map

The contact_map executable identifies residues of the mobile molecule that come within a user-defined distance threshold of the surface across a set of encounter complexes.

```
./contact_map \
  -pdb2          p2_noh.pdb         \
  -complexes     Cu_z_cluster0001_complexes \
  -nb_encounters  5000               \
  -threshold      6.0
```

The expected output is closest_residues.txt, which contains one line per residue listing the encounter-complex indexes for which the residue center of geometry is within the specified distance threshold of the surface.

### Preparation of the LPMO and surface models

The initial structures of NcAA9C (PDB ID: 4D7U)(Borisova *et al.* 2015) and SmAA10A (PDB ID: 8RRY)(Munzone et al. 2024) were retrieved from Protein Data Bank. The structures were protonated at pH = 7.0 using PROPKA(Søndergaard *et al.* 2011) as implemented in PDB2PQR(Dolinsky *et al.* 2004). Amberff19SB(Tian *et al.* 2020) force field was used in LPMOs, and Cu-histidine braces were parameterized using MCPB.PY(Li and Merz 2016) embedded in the Amber24(Case 2025) program package. The starting structures were generated using the GBNeck2 implicit-solvent model (igb=8) with mbondi3 radii using tleap modules in the AMBER24(Case 2025) program package. Prior to the SDA study, a total of 1000 energy minimization steps were performed using steepest descent followed by conjugate gradient minimization with 0.15 M salt concentration to mimic the phycological cellular condition and 1000.0 Å nonbonded cutoff to relax the system.

Cellulose Iβ and its chitin analogue β-chitin were modelled in this study. Cellulose Iβ and β-chitin were generated by Cellulose Builder (Gomes and Skaf 2012) and Chitin Builder, (Malaspina and Faraudo 2024) respectively. The dimensions of the surface models are approximately 200Åx200Å. GLYCAM_06j-1 force field was used to generate pqr files of the surfaces.

### Brownian dynamics simulations – SDA7 protocol

Electrostatic potential grids were calculated by solving the linearized Poisson–Boltzmann equation with APBS (Adaptive Poisson–Boltzmann Solver, version3.0.0)(Gabdoulline and Wade 1996). Solutes were modeled as low-dielectric regions with a dielectric constant of 2, while the surrounding solvent was treated as a high-dielectric continuum with the dielectric constant of 78.4 (dielectric constant of water). The dielectric interface was constructed using the molecular surface definition. The ionic radius and ionic strength were set to 1.5 Å and 50 mM, respectively.

For surfaces, electrostatic potential calculations were carried out using two grid levels defined with a manual multigrid scheme. An initial coarse-grid calculation was performed on a cubic grid of 353×353×353 points with a total grid length of 1059×1059×1059 Å$^3$, corresponding to a grid spacing of approximately 3.0 Å. The grid was centered on the center of geometry of the surface and employed multiple Debye–Hückel boundary conditions. A subsequent focused calculation was carried out using boundary values interpolated from the coarse grid. The focused grid consisted of 353x353x129 points with a uniform grid spacing of 0.75 Å, and was explicitly centered at (100, 100, 0) Å.

For LPMOs, a single grid level was employed using a manually defined multigrid framework. The electrostatic potential calculations were performed on a cubic grid of 200x200x200 Å$^3$ points with a uniform grid spacing of 0.75 Å. The grid was centered on the solute's center of geometry, and the grid boundaries were treated with the multiple Debye–Hückel boundary conditions.

Electrostatic and non-polar desolvation grids were generated using the make_edhdlj_grid module of SDA with a grid spacing of 1 Å. The electrostatic desolvation grids were generated at an ionic strength of 50 mM and by scaling the desolvation forces with a factor of 1.67. Non-polar desolvation forces were scaled by a factor of −0.013.

Translational and rotational diffusion coefficients of the LPMO models were calculated with HYDROPRO (version v10) (I. Torre de, Huertas, and Carrasco 2000, Ortega, Amorós, and García De La Torre 2011) with a radius of the atomic element (AER) of 2.9 Å for proteins. The diffusion coefficients of the surfaces were set to 0.

BD simulations were performed using the Simulation of Diffusional Association (SDA7) software package (version 7.3.5) (Martinez *et al.* 2015). To reduce the number of explicit charges used in the Brownian dynamics simulations, test charges were generated for LPMOs following the standard SDA protocol (Martinez *et al.* 2015), whereas test charges for the surfaces were assigned exclusively to polar atoms (Muñiz-Chicharro, Ganotra, and Wade 2025). In addition, the Effective Charge Model (ECM) implemented in SDA was used to fit the test charges to the electrostatic potential obtained from APBS calculations (Gabdoulline and Wade 1996).

The simulation box extended from (-70, -70, 0) Å to (70, 70, 300) Å to cover the complete surface and to minimize edge effects in periodic boundary conditions (PBCs). A total of 1000 trajectories were run, starting at a box height of 150 Å and terminated when the solute reached the top of the box (at 300 Å) without any time limit restriction (timemax=0). Probe solute and solvent radii were set to 1.77 and 1.4 Å, respectively. A linear variation in the time step was applied, ranging from 1 ps when the center-to-center separation was 60 Å to 20 ps at a separation of 100 Å. Electrostatic interactions between the solute and the surface were calculated in one direction only (oneway_surf_charge = 1) using a corresponding scaling factor of 1 (epfct_oneway_surf = 1.0). Snapshots of the lowest energy were recorded as encounter complexes without using any reaction criteria. Those encounter complexes generated within the same trajectory (ionerun = 1) and with an RMSD < 1.0 Å were merged, and the number of merged encounter complexes were reported in the occupancy column of the encounter complexes file. For each SDA simulation, a total of 60 K encounter complexes were saved.

The software used in this study are as follows and are available for free for academic use: APBS (version 3.0.0),(Jurrus *et al.* 2018) HYDROPRO (version v10), (Ortega, Amorós, and García De La Torre 2011), SDA (version 7.3.5) (Martinez *et al.* 2015)., AMBER24, (Case 2025) PyMOL (version 3.1.8), (Schrödinger LLC, n.d.-b, n.d.-a) and Prot2Surf.

### Contact map analysis

Note that Clusters 1-3 from distance clustering step, that is, Step 1:z-coord, were processed here.

<table>
<tr><td></td><td>Cluster 1</td><td>Cluster 2</td><td>Cluster 3</td></tr>
<tr><td></td><td colspan="3">Example 1: Chitin-active SmAA10A</td></tr>
<tr><td>A</td><td>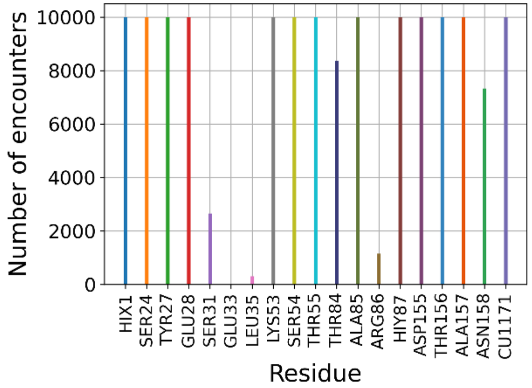
</td><td>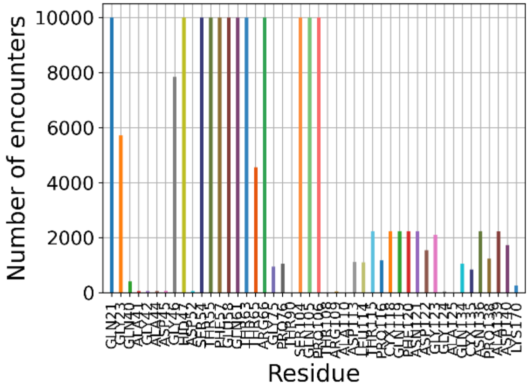
</td><td>N/A</td></tr>
<tr><td>B</td><td>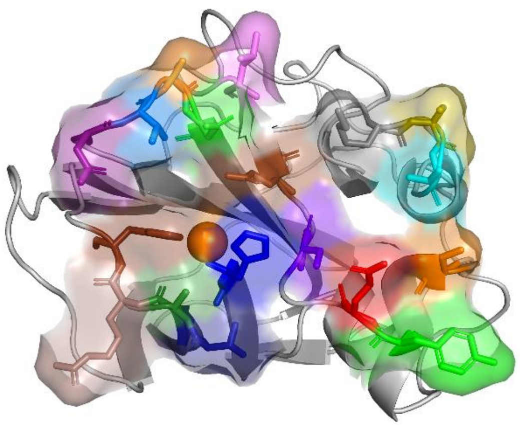</td><td>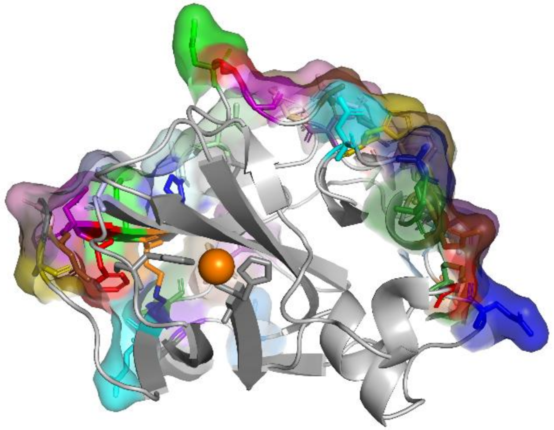</td><td></td></tr>
<tr><td></td><td colspan="3">Example 2: Cellulose-active NcAA9C</td></tr>
</table>

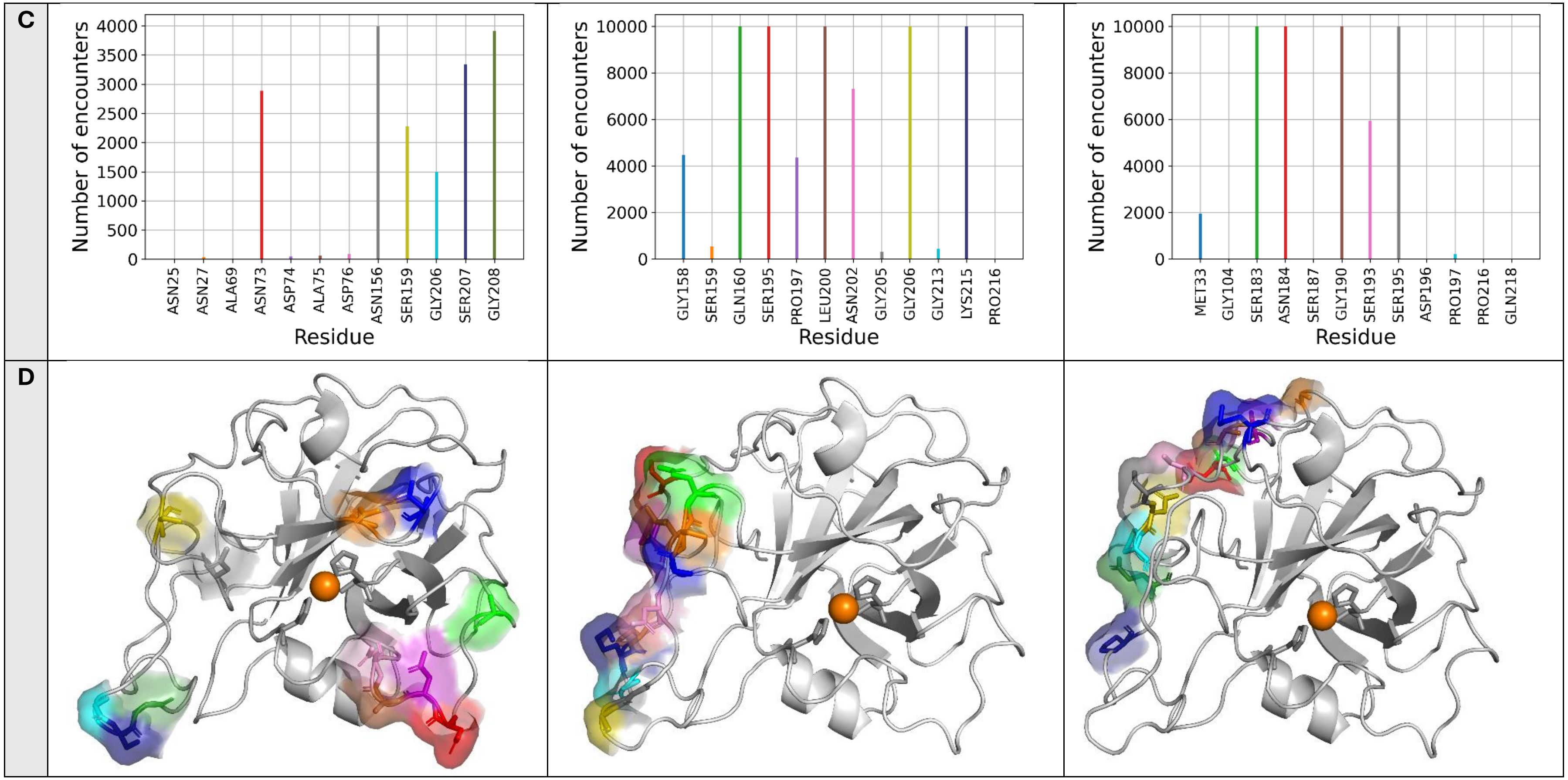

**Figure S4. A, C.** Close contact residues vs number of encounter complexes for the first three clusters of Cu_z distance analysis in Step 1 (if available). **B, D.** The surface and stick presentation of surface contact residues. Contact residues are colored according to the colors of their corresponding vertical lines in A and C. Orange sphere denotes Cu ion. For visual clarification, surfaces were not illustrated here.

### Raw output data

**Table S1.** The Prot2Surf analysis output of the example cases.

#### Example cases 1: Chitin active SmAA10A

***Cu_z_clust_info.txt***

| Clust | pop. | Repr. | Average | SD |
|---|---|---|---|---|
| Cluster1 | 19086 | 1 | 5.096 | 0.732 |
| Cluster2 | 40914 | 41235 | 20.879 | 0.485 |

***2D_angle_thres7.5_clust_info.txt***

| Clust | pop. | Repr. | Average | SD |
|---|---|---|---|---|
| Cluster1 | 9238 | 5108 | 5.581 | 4.028 |
| Cluster2 | 8118 | 13324 | 106.685 | 2.872 |

***regioselectivity_occ.log***

| | | |
|---|---|---|
| C1_dist_thres7.5_complexes: | 7931/4383366 | (0.18%) |
| C4_dist_thres7.5_complexes: | 679713/4383366 | (15.51%) |
| C1_C4_dist_thres7.5_complexes | 850833/4383366 | (19.41%) |
| No reaction: | 2844889/4383366 | (64.90%) |

**results_closest_residues – contact map**

***Cu_z_cl1.dat***

| #ResidueNumber | ResidueName | Occurrences |
|---|---|---|
| 1 | HIX1 | 10000 |
| 24 | SER24 | 10000 |
| 27 | TYR27 | 10000 |
| 28 | GLU28 | 10000 |

#### Example cases 2: Cellulose active NcAA9C

***Cu_z_clust_info.txt***

| Clust | pop. | Repr. | Average | SD |
|---|---|---|---|---|
| Cluster1 | 3992 | 1890 | 7.142 | 0.201 |
| Cluster2 | 43536 | 24117 | 22.180 | 0.737 |
| Cluster3 | 11566 | 53224 | 25.610 | 0.245 |
| Cluster4 | 3 | 59095 | 28.418 | 0.047 |
| Cluster5 | 903 | 59522 | 35.763 | 0.307 |

***2D_angle_thres7.5_clust_info.txt***

| Clust | pop. | Repr. | Average | SD |
|---|---|---|---|---|
| Cluster1 | 89 | 43 | 16.668 | 4.460 |
| Cluster2 | 178 | 167 | 37.493 | 6.094 |
| Cluster3 | 47 | 294 | 59.878 | 3.006 |
| Cluster4 | 325 | 486 | 78.724 | 5.431 |
| Cluster5 | 294 | 802 | 100.070 | 8.089 |
| Cluster6 | 467 | 1182 | 148.274 | 9.97 |

***regioselectivity_occ.log***

| | | |
|---|---|---|
| C1_dist_thres7.5_complexes: | 4894/1513211 | (0.32%) |
| C4_dist_thres7.5_complexes: | 7827/1513211 | (0.52%) |
| C1_C4_dist_thres7.5_complexes | 4771/1513211 | (0.32%) |
| No reaction: | 1495719/1513211 | (98.84%) |

***Cu_z_cl1.dat***

| #ResidueNumber | ResidueName | Occurrences |
|---|---|---|
| 25 | ASN25 | 8 |
| 27 | ASN27 | 39 |
| 69 | ALA69 | 10 |
| 73 | ASN73 | 2887 |

| | | | | | |
|---|---|---|---|---|---|
| 31 | SER31 | 2650 | 74 | ASP74 | 48 |
| 33 | GLU33 | 10 | 75 | ALA75 | 65 |
| 35 | LEU35 | 308 | 76 | ASP76 | 96 |
| 53 | LYS53 | 10000 | 156 | ASN156 | 3990 |
| 54 | SER54 | 10000 | 159 | SER159 | 2280 |
| 55 | THR55 | 10000 | 206 | GLY206 | 1496 |
| 84 | THR84 | 8377 | 207 | SER207 | 3343 |
| 85 | ALA85 | 10000 | 208 | GLY208 | 3908 |
| 86 | ARG86 | 1156 | | | |
| 87 | HIY87 | 10000 | | | |
| 155 | ASP155 | 10000 | | | |
| 156 | THR156 | 10000 | | | |
| 157 | ALA157 | 10000 | | | |
| 158 | ASN158 | 7333 | | | |
| 171 | CU1171 | 10000 | | | |
| **Cu_z_cl2.dat** | | | **Cu_z_cl2.dat** | | |
| **#ResidueNumber** | **ResidueName** | **Occurrences** | **#ResidueNumber** | **ResidueName** | **Occurrences** |
| 21 | GLN21 | 10000 | 158 | GLY158 | 4483 |
| 23 | GLY23 | 5718 | 159 | SER159 | 542 |
| 40 | GLN40 | 413 | 160 | GLN160 | 10000 |
| 41 | ALA41 | 60 | 195 | SER195 | 10000 |
| 42 | GLY42 | 59 | 197 | PRO197 | 4362 |
| 44 | ALA44 | 60 | 200 | LEU200 | 10000 |
| 45 | ASP45 | 68 | 202 | ASN202 | 7316 |
| 46 | GLY46 | 7846 | 205 | GLY205 | 321 |
| 47 | HID47 | 10000 | 206 | GLY206 | 10000 |
| 52 | ASP52 | 60 | 213 | GLY213 | 437 |
| 54 | SER54 | 10000 | 215 | LYS215 | 10000 |
| 55 | THR55 | 10000 | 216 | PRO216 | 8 |
| 57 | PHE57 | 10000 | | | |
| 58 | GLU58 | 10000 | | | |

| | | |
|---|---|---|
| 61 | GLN61 | 10000 |
| 63 | THR63 | 10000 |
| 65 | THR65 | 4561 |
| 66 | ARG66 | 10000 |
| 75 | GLY75 | 945 |
| 76 | PRO76 | 1062 |
| 90 | THR90 | 3 |
| 104 | SER104 | 10000 |
| 105 | GLN105 | 10000 |
| 106 | PRO106 | 10000 |
| 108 | THR108 | 24 |
| 109 | ARG109 | 55 |
| 110 | ALA110 | 14 |
| 113 | ASP113 | 1112 |
| 114 | LEU114 | 1100 |
| 115 | THR115 | 2237 |
| 116 | PRO116 | 1176 |
| 118 | CYX118 | 2236 |
| 119 | GLN119 | 2237 |
| 120 | PHE120 | 2237 |
| 121 | ASN121 | 2237 |
| 122 | ASP122 | 1552 |
| 123 | GLY123 | 2106 |
| 124 | GLY124 | 54 |
| 125 | ALA125 | 2 |
| 134 | GLN134 | 1062 |
| 135 | CYX135 | 844 |
| 136 | ASN136 | 2237 |
| 138 | PRO138 | 1237 |
| 139 | ALA139 | 2225 |
| 140 | ASP140 | 1735 |

| | |
|---|---|
| 170 LYS170 262 | |
| **Cu_z_cl3.dat** | **Cu_z_cl3.dat** |
| N/A | **#ResidueNumber ResidueName Occurrences**<br>33 MET33 1957<br>104 GLY104 6<br>183 SER183 10000<br>184 ASN184 10000<br>187 SER187 6<br>190 GLY190 10000<br>193 SER193 5945<br>195 SER195 10000<br>196 ASP196 8<br>197 PRO197 211<br>216 PRO216 1<br>218 GLN218 10 |
| **Cu_z_cl4.dat** | **Cu_z_cl4.dat** |
| N/A | **#ResidueNumber ResidueName Occurrences**<br>183 SER183 3<br>184 ASN184 3<br>215 LYS215 3<br>216 PRO216 3 |
| **Cu_z_cl5.dat** | **Cu_z_cl5.dat** |
| | **#ResidueNumber ResidueName Occurrences**<br>47 SER47 519<br>51 THR51 113<br>95 ASP95 902<br>96 ASN96 903<br>142 PRO142 903<br>143 GLY143 884<br>144 GLN144 2<br>173 ASN173 437 |

| | | |
|---|---|---|
| | 175 THR175 | 903 |
| | 176 GLY176 | 903 |
| | 177 GLY177 | 693 |
| | 178 GLY178 | 725 |
| | 179 SER179 | 56 |

### References

Borisova AS, Isaksen T, Dimarogona M *et al.* Structural and functional characterization of a lytic polysaccharide monooxygenase with broad substrate specificity. *Journal of Biological Chemistry* 2015;**290**(38):22955–69. https://doi.org/10.1074/jbc.M115.660183.

Case DA; AHM; BSIY; BJT; BSR; CFS; et al. Amber 2024. *University of California, San Francisco* 2025.

Dolinsky TJ, Nielsen JE, McCammon JA *et al.* PDB2PQR: an automated pipeline for the setup of Poisson–Boltzmann electrostatics calculations. *Nucleic Acids Res* 2004;**32**(suppl_2):W665–7. https://doi.org/10.1093/NAR/GKH381.

Gabdoulline RR, Wade RC. Effective charges for macromolecules in Solvent. *J Phys Chem* 1996;**100**:3868–78.

Gomes TCF, Skaf MS. Cellulose-Builder: A toolkit for building crystalline structures of cellulose. *J Comput Chem* 2012;**33**(14):1338–46. https://doi.org/10.1002/jcc.22959.

Jurrus E, Engel D, Star K *et al.* Improvements to the APBS biomolecular solvation software suite. *Protein Science* 2018;**27**(1):112–28. https://doi.org/10.1002/pro.3280.

l. Torre JG de, Huertas M, Carrasco B. Calculation of hydrodynamic properties of globular proteins from their atomic-level structure. *Biophys J* 2000;**78**(2):719–30. https://doi.org/10.1016%2FS0006-3495(00)76630-6.

Li P, Merz KM. MCPB.py: A Python Based Metal Center Parameter Builder. *J Chem Inf Model* 2016;**56**(4):599–604. https://doi.org/10.1021/acs.jcim.5b00674.

Malaspina D, Faraudo J. Chitin Builder: a VMD tool for the generation of structures of chitin molecular crystals for atomistic simulations. *J Open Source Softw* 2024;**9**(93):5771. https://doi.org/10.21105/joss.05771.

Martinez M, Bruce NJ, Romanowska J *et al.* SDA 7: A modular and parallel implementation of the simulation of diffusional association software. *J Comput Chem* 2015;**36**(21):1631–45. https://doi.org/10.1002/jcc.23971.

Muñiz-Chicharro A, Ganotra GK, Wade RC. A Multiscale Simulation Approach to Compute Protein–Ligand Association Rate Constants by Combining Brownian Dynamics and Molecular Dynamics. *J Chem Inf Model* 2025;**65**(20):11215–31. https://doi.org/10.1021/acs.jcim.5c01488.

Munzone A, Pujol M, Tamhankar A *et al.* Integrated Experimental and Theoretical Investigation of Copper Active Site Properties of a Lytic Polysaccharide Monooxygenase from Serratia marcescens. *Inorg Chem* 2024;**63**(24):11063–78. https://doi.org/10.1021/acs.inorgchem.4c00602.

Ortega A, Amorós D, García De La Torre J. Prediction of Hydrodynamic and Other Solution Properties of Rigid Proteins from Atomic- and Residue-Level Models. *Biophys J* 2011;**101**(4):892–8. https://doi.org/10.1016/J.BPJ.2011.06.046.

Schrödinger LLC. *The AxPyMOL Molecular Graphics Plugin for Microsoft PowerPoint, Version 3.1.8*. n.d.

Schrödinger LLC. *The PyMOL Molecular Graphics System, Version 3.1.8*. n.d.

Søndergaard CR, Olsson MHM, Rostkowski M *et al.* Improved Treatment of Ligands and Coupling Effects in Empirical Calculation and Rationalization of pKa Values. *J Chem Theory Comput* 2011;**7**(7):2284–95. https://doi.org/10.1021/CT200133Y.

Tian C, Kasavajhala K, Belfon KAA *et al.* Ff19SB: Amino-Acid-Specific Protein Backbone Parameters Trained against Quantum Mechanics Energy Surfaces in Solution. *J Chem Theory Comput* 2020;**16**(1):528–52. https://doi.org/10.1021/ACS.JCTC.9B00591/SUPPL_FILE/CT9B00591_SI_002.ZIP.
